# Effect of Infection with, and Treatment of, Sensitive and Resistant Strains of *Teladorsagia Circumcincta* on the Ovine Intestinal Microbiota

**DOI:** 10.1101/304386

**Authors:** Craig A. Watkins, Dave J. Bartley, Burcu Gündüz Ergün, Büşra Yıldızhan, Tracy Ross-Watt, Alison A. Morrison, Maria J. Rosales Sanmartín, Fiona Strathdee, Leigh Andrews, Andrew Free

## Abstract

Nematodes are one of the main impactors on health, welfare and productivity of farmed animals. *Teladorsagia circumcincta* is arguably one of the most globally important nematode species in sheep. Control of these nematode infections is essential and heavily reliant on chemotherapy (anthelmintics), but this has been complicated by the development of anthelmintic resistance. In mammals the composition of the intestinal microbiota has been shown to have a significant effect on overall health. The interactions between host, microbiota and pathogens are complex and influenced by numerous factors. In this study, the interactions between *T. circumcincta* infections and microbial composition and abundance were investigated. In a preliminary study the intra-and inter-individual diversity and composition of the microbiota of grazing sheep was assessed in two distinct regions of the ovine intestinal tract, the terminal ileum (TI) and rectal contents. Additionally, the effects of experimental infection of sheep with two strains of *T. circumcincta* (anthelmintic resistant or sensitive) on the microbiota were assessed with and without anthelmintic (monepantel) administration. The inter-animal variability was greater in the TI compared to the rectal samples. However, the alpha-diversity (species richness) was significantly lower in the TI samples. In the experimental study, clear differences were observed between successfully treated animals and those sheep that were left untreated and/or those carrying resistant nematodes. Differences in microbiota between the four different experimental conditions were observed and potential predictive biomarkers were identified. In particular, a restoration of potentially beneficial *Bifidobacteria* sp. in successfully-treated animals was observed.

## IMPORTANCE

Roundworms are one of the main impactors on health, welfare and productivity of farmed animals. The roundworm *Teladorsagia circumcincta* is arguably one of the most globally important in sheep. Control of these roundworm infections is essential and heavily reliant on chemotherapy, but this has been complicated by the development of drug resistance. In mammals including humans, the composition of the intestinal microbiota has shown significant effects on health. Interactions between roundworm infections and microbial composition and abundance were investigated. The significance of this current work is that: i) the within-and between-animal variability in microbial composition was assessed. ii) the interaction between roundworm and microbiota of sheep with and without drug administration were evaluated. The use of resistant and sensitive helminth strains was compared. iii) successful removal of pathogenic roundworms results in a higher abundance of specific microflora indicating those organisms that are associated with a healthy gut microflora.

## INTRODUCTION

Within the gastrointestinal tract of mammals including humans, micro-organisms form a complex relationship with their hosts, be they commensal symbionts, pathobionts or pathogenic, as a consequence of co-evolution to provide homeostasis in the host intestinal tract (1–3). In healthy animals, the microbiota provides essential nutrients and protection against the colonisation by pathogenic species (4). However, constant regulation is required to prevent breakdown of these essential relationships between microorganisms and mammals. Where commensal bacteria, invading pathogenic bacteria and/or opportunistic pathobionts colonise similar ecological niches, competition for available nutrients occurs (5). Such interactions can lead to perturbations in the microbial communities leading to dysfunction of the gastro-intestinal tract (6). The response of commensal microbiota therefore needs to be robust to out-compete and deny the incoming, infectious agent from colonising and proliferating (7, 8). However, when the microbiota becomes disrupted, unstable or damaged (for example due to antibiotics, pathogenic infections, physiological and/or environmental stresses), the ability of the commensal microorganisms to maintain a competent resistance to colonisation by pathogenic organisms can be compromised. These invading pathogenic microorganisms have their own arsenal to counteract the defensive mechanisms developed by the commensal microflora and the host, including physiological and immunological responses (5, 9–12). Gastrointestinal parasites provide just the type of disruption that results in breach of physical boundaries, inflammation and/or modulation of the immune response of the host, offering an opportunity for invading microorganisms to colonise (13). A number of factors exist which impact on the helminth-host microbiota interaction including; age, sex, nutrition and immune status (14, 15). Together, the host, its microbial community and the presence of parasites are known to shape both the microbial landscape and host health (3).

The impact of gastro-intestinal parasites on the microbiota has not been extensively studied in mammals, let alone specifically in sheep and other ruminants (3). However, it is clear that the perturbation of the gastrointestinal tract can be pronounced when pathogenic micro-organisms are introduced, either naturally or experimentally (4, 13, 16, 17). *Teladorsagia circumcincta* is a pathogenic parasitic nematode that can cause severe gastroenteritis. It is one of the most common gastrointestinal nematodes in sheep worldwide within the temperate zone (18), causing considerable morbidity and occasionally mortality in heavily infected animals (19). Clinical symptoms are attributed to both excretory-secretory (ES) products derived from the invading nematode and to histopathological tissue damage (20, 21). Control of *T. circumcincta* remains a challenge, with anthelmintics being the mainstay of control although the use of some treatments has become restricted due to increased, and widespread, multi-drug resistance developed by *T. circumcincta* (22, 23).

The pathogenic nature of gastro-intestinal round worms has a significant effect on the microbiota of the gut (3). However, quantifying precisely the dynamics of the microbial community has not been possible until the advent of culture-independent 16S rDNA-based sequence analysis and whole genome shotgun high-throughput sequencing (24). In this study, Illumina MiSeq technology was used to sequence amplicons of the V4 region of the 16S rRNA gene, to determine the inherent variability of the gut microbiota between samples obtained from two different areas of the intestinal tract (rectum and terminal ileum) in six identically-treated sheep. The same approach was then used to define the effect of infection with, and treatment of, sensitive and resistant strains of *T. circumcincta* on the ovine intestinal microbiota, via rectal faecal (RF) composition.

## RESULTS

### Inherent variation of the ileal and rectal microbiotas

In this initial study, samples from two distinct regions of the gut, the terminal ileal contents (TIC) and rectal faeces (RF), in six identically managed sheep (TIC 1-6 and RF 1-6 respectively), were analysed to investigate intra-and inter-sheep variation of the bacterial communities. Good’s coverage estimates on sequence data obtained from these 12 samples exceeded 99% for all samples, demonstrating that the sequencing depth was adequate to capture true diversity (minimum ~119,000 sequences reads per sample; Table S1). Consistent with alpha rarefaction curves of operational taxonomic unit (OTU) richness (Fig. 1A), Shannon diversity (mean ± SD at maximum rarefaction depth) was lower in the TIC samples (5.054 ± 1.595) than the RF samples (7.869 ± 0.252), with a corresponding reduction in Pielou’s evenness (0.668 ± 0.262 compared to 0.951 ± 0.048). This shows that there is a significantly lower diversity (Welch’s t-test, p = 0.0006) of the microbial communities found in the TIC compared to the rectum (Fig. 1A), with the former dominated by a few abundant OTUs. However, alpha diversity was also more variable in the TIC samples than in the RF samples, as illustrated by the error bars in Fig. 1A. Consistent with this, compositional variability, shown by Principal Coordinates Analysis (PCoA) ordination based on Bray-Curtis similarity (S), was greater between the TIC samples (mean S = 38.3; SEM= 6.21) than between those taken from the rectum (Fig. 1B). The RF samples showed greater similarity in composition (mean S = 60.9; SEM = 1.47) than the TIC samples, a difference shown to be highly significant by PERMDISP analysis of multivariate dispersions (F = 12.89; P = 0.003). As illustrated by the PCoA plot, the tightly-clustered RF samples were also compositionally distinct from the more variable TIC samples, verified by a PERMANOVA test (Pseudo-F = 13.62; P =0.004). Vectors illustrated in Fig. 1B show that the abundances of the major phyla *Bacteroidetes, Firmicutes* and *Proteobacteria* were all closely correlated (r > 0.9) with the axes of the plot. Elevated abundances of *Bacteroidetes* were associated with the RF samples, while *Proteobacteria* and *Firmicutes* were associated with the TIC, but in proportions which vary between the individual animals.

**FIG 1.**
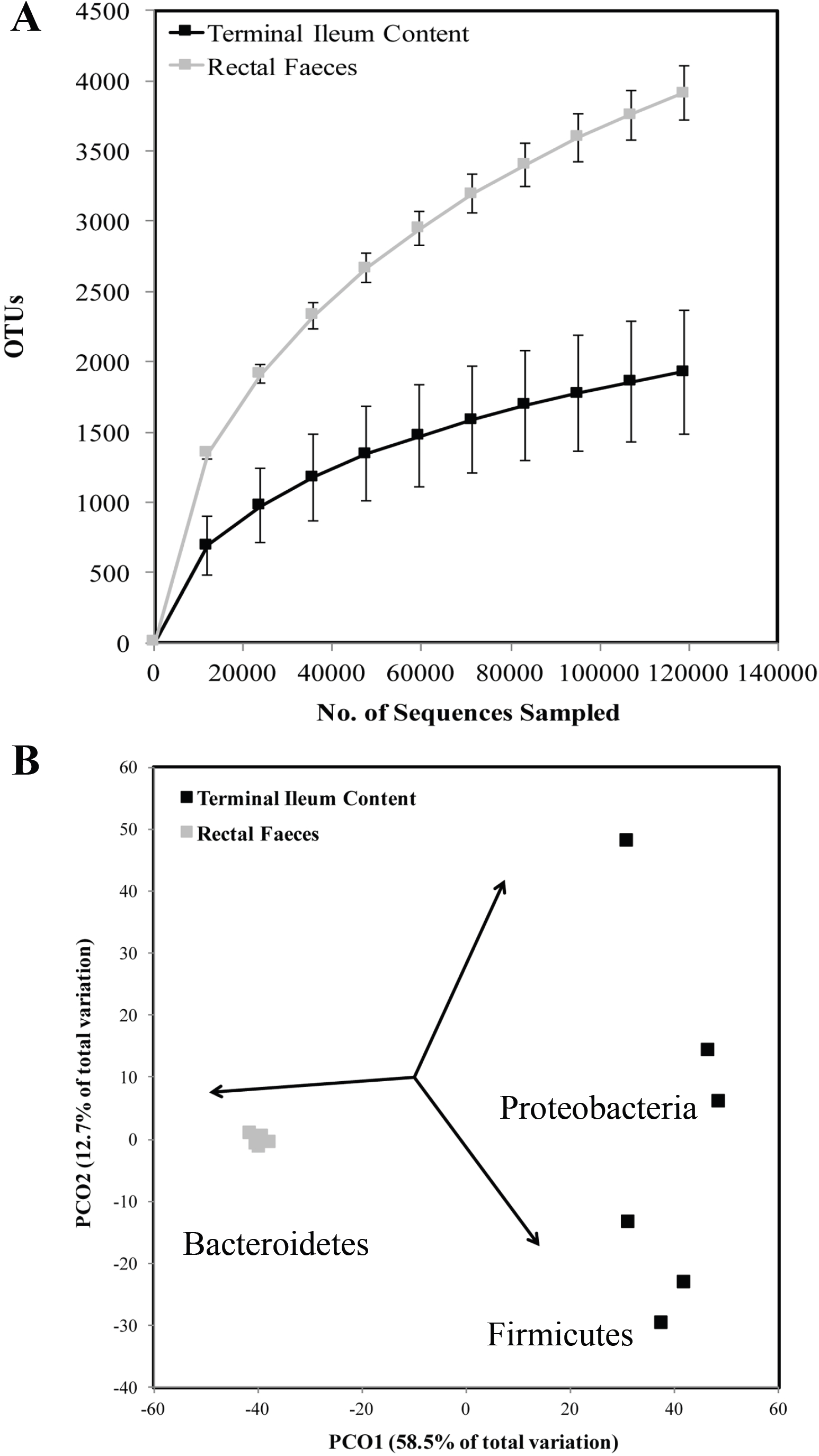
(A) Alpha rarefaction curves for OTUs in terminal ileal content (TIC) and rectal faecal (RF) samples (n=6 for each sample type). Error bars represent standard deviations. (B) PCoA ordination based on Bray-Curtis similarity of OTU distributions for TIC and RF samples. Vectors show Pearson correlations between the abundances of major Phyla and the PCoA axes.

Relative abundance plots of bacterial Orders showed the different taxonomic distribution patterns between TIC and RF samples (Fig. S1). Twelve different Orders of bacteria were identified in the TC and RF samples at a limit of >4% abundance. *Clostridiales* was the dominant order for TIC samples (35-70%) except TIC-5 (18%), where *Enterobacteriales* was the dominant population of bacteria (76% of the total). For RF samples *Clostridiales* (34-43%) and *Bacteroidales* (38-50%) made similar contributions to the bacterial abundance. Average relative abundance (mean ± SD) of *Enterobacteriales* was generally higher in TIC samples (12.5 ± 6.77%, TIC-5 excluded) than RF samples (0.64 ± 0.55%). The relative abundance and distribution of the five most abundant bacterial OTUs in each of the TIC and RF samples was analysed and illustrated using a heatmap (Fig. 2), indicating the very different profiles of microbiota obtained from the two different sampling sites. This analysis suggests that OTUs from the Genus *Ruminococcaceae* UCG-005 (25), various *Bacteroidales* OTUs, and OTUs from the Genera *Campylobacter* and *Akkermansia* were over-represented in the RF samples compared to the ileal samples. In contrast, OTUs corresponding to various Genera from the Order *Clostridiales*, the species *Escherichia coli*, the Genus *Turicibacter*, the Order *Victivallales*, the Family *Bifidobacteriaceae* and the Genus *Methanobrevibacter* were better represented in the TIC samples. A single OTU (denovo19121) assigned to the genus “*Candidatus Hepatincola*” (26) was abundant in one sheep TIC sample (TIC5), but poorly represented in all other samples. Full details of the percentage abundance and taxonomy of the 15 most abundant OTUs in each sample, including those shown in Fig. 2, are presented in Table S2.

**FIG 2.**
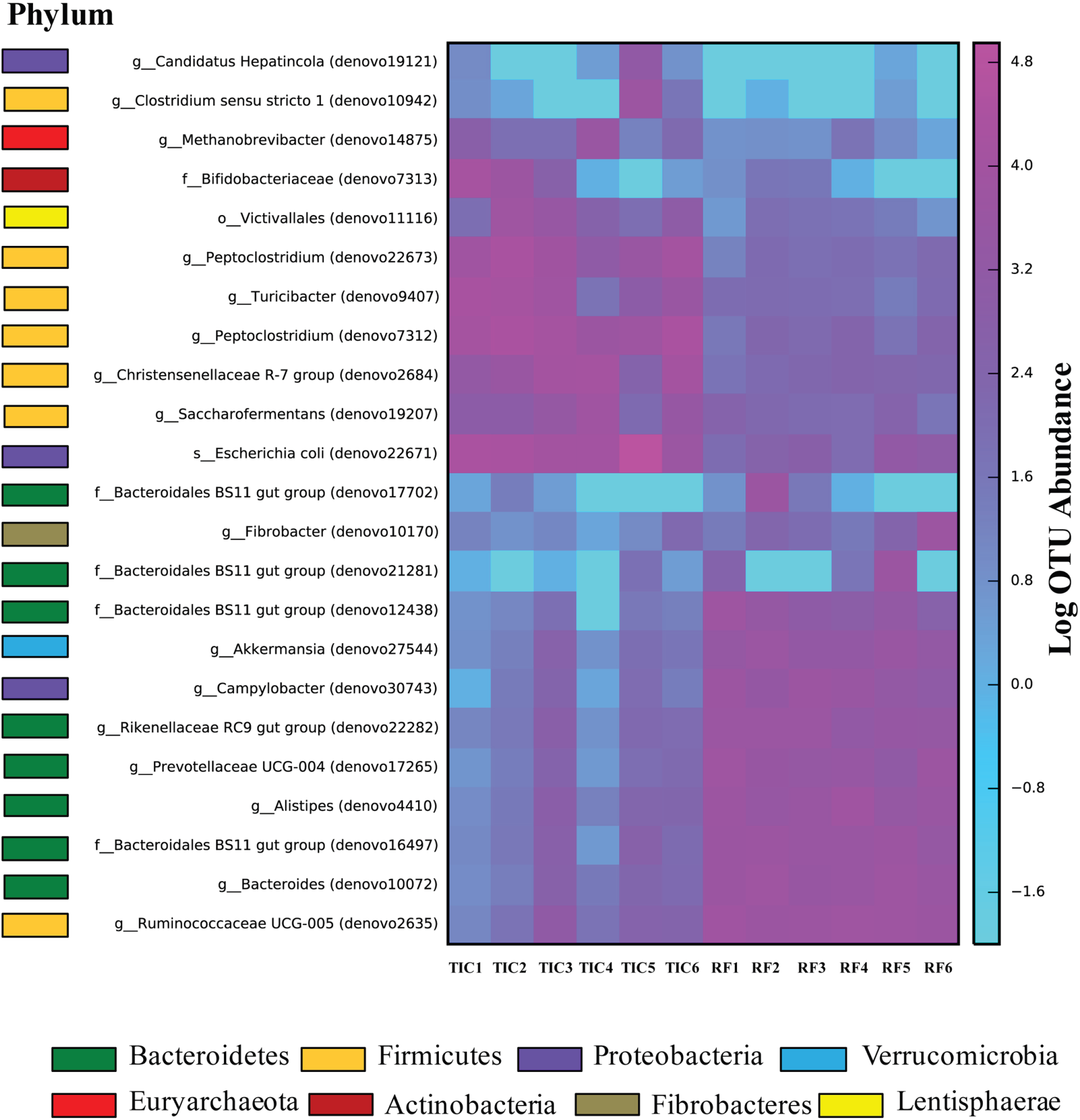
Heat map representing the abundances of individual OTUs in the TIC and RF samples. The five most abundant OTUs in each sample were included in the analysis. The most accurate levels of taxonomic classification for each OTU is shown next to the OTU identifier, and Phylum membership is indicated with coloured bars.

These experiments showed that the inherent variability of the microbiota between sheep was much less for RF samples than for TIC samples. Therefore, RF samples were collected to investigate the effect of treatment for sensitive versus resistant *T. circumcincta* on microbiota composition.

### Effect of infection with, and treatment of, sensitive and resistant strains of *T. circumcincta*

Sequence data were obtained on the RF samples from four groups of five sheep (n = 20) infected with sensitive or resistant strains of *T. circumcincta* and either treated with monepantel (MPTL) or left untreated (Table 1). Good’s coverage estimates were ≥ 97% for all samples, demonstrating that the sequencing depth was adequate to capture true diversity (minimum ~118,000 sequences reads per sample; Table S1). Observed species rarefaction curves for the four groups of infected sheep (susceptible treated, ST; susceptible untreated, SUT; resistant treated, RT; resistant untreated, RUT) showed high diversity within these faecal samples (Fig. 3A), consistent with the faecal samples analysed in the first experiment. However, Shannon diversities (mean ± SD at maximum rarefaction depth) for those sheep left untreated (SUT, RUT) and those not successfully treated (RT) for worm infections were higher (8.694 ± 0.229) than those for samples representing sheep that had been successfully treated (ST) (8.434 ± 0.192), a difference which, though small, is significant at the 5% level (Welch’s t-test, p = 0.037). This suggests that there is a slight but detectable increase in microbial species diversity in sheep with active nematode infections in this experiment.

**TABLE 1.** *Teladorsagia circumcincta* isolate designations pre-and post-selection for monepantel resistance

| Original isolate designation | Anthelmintic sensitivity<sup>a</sup> |  |  |  | Anthelmintic administration (dose rate; mg kg<sup>-1</sup> body weight) | Designation during microbiota characterisation |
| --- | --- | --- | --- | --- | --- | --- |
|  | BZ | LV | ML | MP |  |  |
| MTci7 | R | R | R | S | None | SUT |
|  |  |  |  |  | Monepantel - Zolvix® (2.5) | ST |
| MTci7-12 | R | R | R | R | None | RUT |
|  |  |  |  |  | Monepantel - Zolvix® (2.5) | RT |

<sup>a</sup>BZ, benzimidazole; LV, levamisole; ML, macrocyclic lactone (ivermectin and milbemycin); MP, monepantel; S, sensitive to anthelmintic class; R, resistant to anthelmintic class

**FIG 3.**
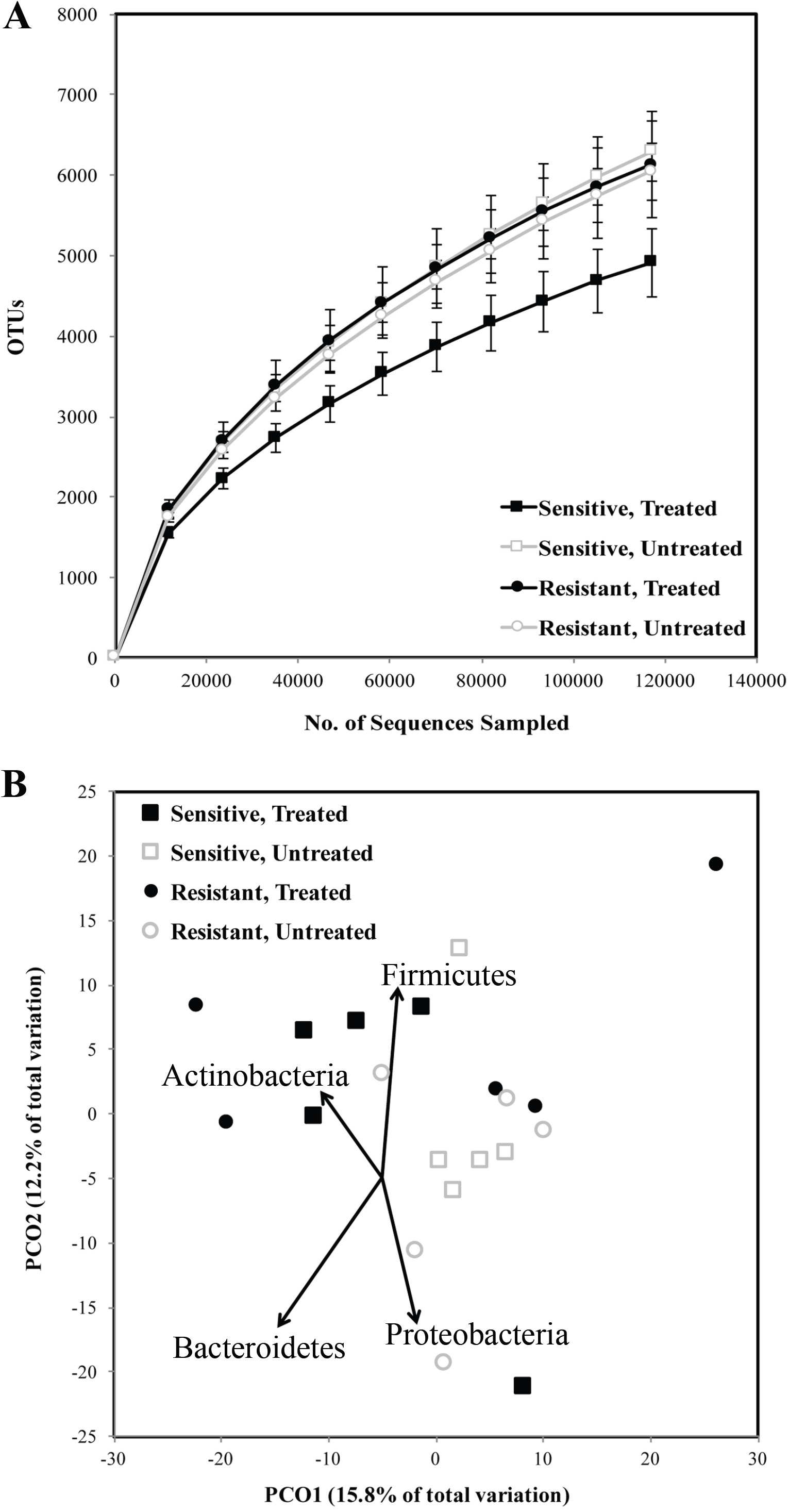
(A) Alpha rarefaction curves for OTUs in RF samples from monepantel-treated and untreated sheep infected with sensitive or resistant *T. circumcincta* (n=5 for each treatment). Error bars represent standard deviations. (B) PCoA ordination based on Bray-Curtis similarity of OTUs in RF samples from monepantel treated and untreated sheep infected with sensitive or resistant *T. circumcincta*. Vectors show Pearson correlations between the abundances of major Phyla and the PCoA axes.

At the Order level of taxonomy, the composition of the RF samples from all four treatment groups were dominated by *Bacteroidales* (30-49%) and *Clostridiales* (27-48%; Fig. S2). There were also significant but smaller abundances of *Spirochaetales* (1-4%) and *Campylobacterales* (0.4-6%). To investigate finer-scale similarity between the four different groups, Bray-Curtis similarity based on OTU abundances was visualised by PCoA ordination (Fig. 3B). This analysis revealed a large degree (72%) of inter-individual variability unrepresented in the first two PCoA axes, which nevertheless suggested some compositional differences associated with the four treatment groups, and showed significant correlations with the abundances of the phyla *Bacteroidetes, Firmicutes* and *Proteobacteria* (P ≤ 0.003) and *Actinobacteria* (P = 0.023). Despite the observed variability, those sheep groups with ongoing nematode infections (SUT, RT, RUT) showed a significant difference in composition from those treated successfully (ST; PERMANOVA Pseudo-F = 1.594, P = 0.016). No such statistical significance was detected for the factors of drug treatment or *T. circumcincta* strain (P > 0.18). To isolate the variation due to the pre-defined treatment groups in the experimental set-up, a Canonical Analysis of Principal coordinates (CAP) ordination plot based on these groups was used. The CAP plot (using the same OTU-level similarity data as in Fig. 3B) showed that there was a clear separation of the successfully-treated group (ST) from the untreated or resistant groups (SUT, RT, RUT) on the primary CAP1 axis, while the three latter groups are separated from each other on the CAP2 axis (Fig. 4A). These treatment group-specific separations are associated with high eigenvalues (0.939 and 0.672 for CAP1 and CAP2 respectively), and a significant first canonical correlation (P = 0.005) (27). As expected, the vectors in Fig. 4A show that there is a strong correlation (r = 0.626; P = 0.002) between worm burden and the sample groups as separated on the CAP1 axis. There is also a strong positive correlation between the abundance of OTUs assigned to the Phylum *Actinobacteria* and the ST group (r = 0.651; P = 0.001), while the separation of the other three treatment groups on the CAP2 axis is correlated with the abundance of the Phylum *Proteobacteria* (r = 0.805; P < 0.0001), although driven mainly by a single RUT sample (Fig. S2).

**FIG 4.**
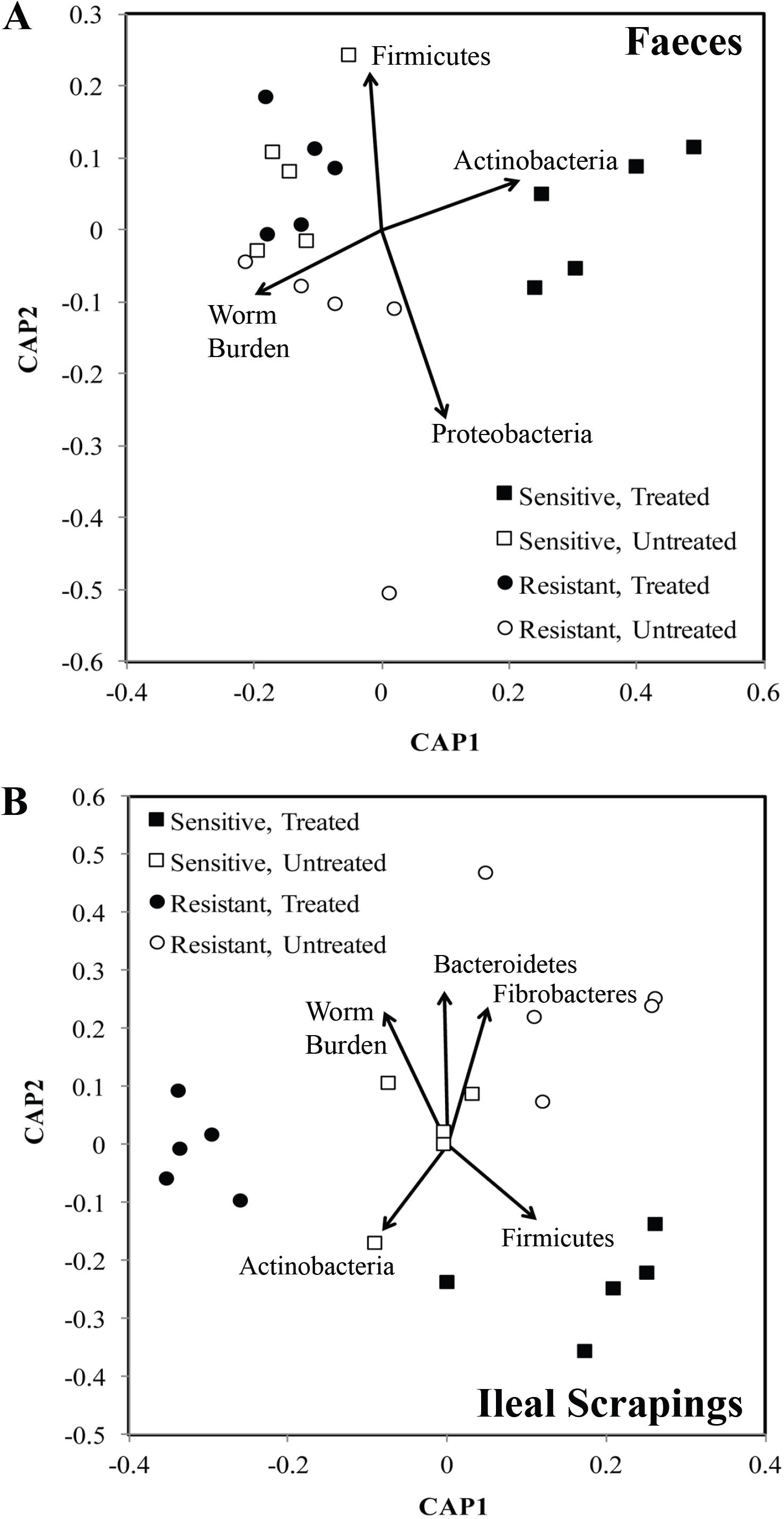
CAP ordination based on Bray-Curtis similarity of OTUs in RF samples (A) and TIMS (B) from monepantel-treated and untreated sheep, infected with sensitive or resistant *T. circumcincta* (n=5 for each treatment). Vectors show Pearson correlations between the abundances of major Phyla or the worm burden and the CAP axes. Shaded data points represent anthelmintic treatment, while squares and circles indicate the sensitive and resistant *T. circumcincta* strains respectively.

To confirm the effects detected in faecal samples, we analysed microbial DNA present in terminal ileal mucosal scraped (TIMS) samples taken from the same animals at post mortem. As the yield of microbial DNA from these samples was much lower than from faecal pellets or TIC, lower numbers of sequences per sample were obtained (Table S1). However, rarefaction curves (Fig. S3A) and Good’s coverage estimates (>94% for all samples) suggested that the low microbial diversity in these samples was covered adequately at this sequencing depth. OTU richness and Shannon diversity were relatively low in these samples, consistent with our previous analysis of the terminal ileal contents (TIC). However, there was a small but significant (Welch’s t-test; p = 0.015) decrease in diversity in those samples from sheep treated with monepantel compared to the untreated sheep: Shannon diversities (mean ± SD) were 5.934 ± 1.039 in treated animals compared to 6.972 ± 0.480 in untreated animals.

Sixteen different Orders of bacteria were identified in the TIMS samples at a limit of >4% abundance (Fig. S3B), including sequences corresponding to the 16S rRNA gene from the remnant chloroplast (apicoplast) of the apicomplexan parasite *Eimeria*. The most abundant Order across the sample set was *Clostridiales* (12-64%). Although large inter-sample variability was observed, similar to the TIC microbiota samples, a PERMANOVA test showed that there was a significant compositional difference between successfully treated and nematode-infected sheep (Pseudo-F = 1.073; P = 0.027). A CAP analysis of Bray-Curtis similarities based on OTU abundances (Fig. 4B) indicated that, as for the faecal samples, the sheep with successfully-treated nematode infections clustered separately from those with untreated or resistant infections (eigenvalues = 0.943 and 0.881; trace statistic P = 0.008). There were also strong correlations between the CAP axes and worm burden (r = 0.696; P = 0.0003), as expected, and the abundance of *Bacteroidetes* (r = 0.753; P < 0.0001) and *Fibrobacteres* (r = 0.696; P = 0.0003), with samples derived from the untreated sheep (SUT and RUT). Weaker but significant correlations were also observed with the abundance of *Firmicutes* (r = 0.491; P = 0.014) and *Actinobacteria* (r = 0.486; P = 0.015) with ST samples (Fig. 4B).

### Identification of biomarkers defining infected and uninfected sheep

To evaluate structural differences in the constituent microbial communities of infected and uninfected sheep, a heatmap illustrating the distribution of the five most abundant OTUs in the RF samples was constructed (Fig. 5). The vast majority of these OTUs were assigned to taxa within the Order *Bacteroidales*, with individual OTUs assigned to the genera *Ruminococcaceae* UCG-005, *Campylobacter, Helicobacter* and *Methanocorpusculum* also represented. Full details of the percentage abundance and taxonomy of the 15 most abundant OTUs in each sample, including those shown in Fig. 5, is presented in Table S3. Although some minor differences between individual samples at the OTU level were apparent, differences specific to the four treatment groups were not obvious among this subset of abundant OTUs. The TIMS samples from the same sheep exhibited much more variability in the abundant OTUs between animals, and a preponderance of OTUs from the Phylum *Firmicutes*. Variation in most of these OTUs between the four treatment groups was not readily apparent, but a single *Bifidobacterium* OTU (denovo3328) was largely restricted to the successfully-treated ST group (Fig. S4; Table S4).

**FIG 5.**
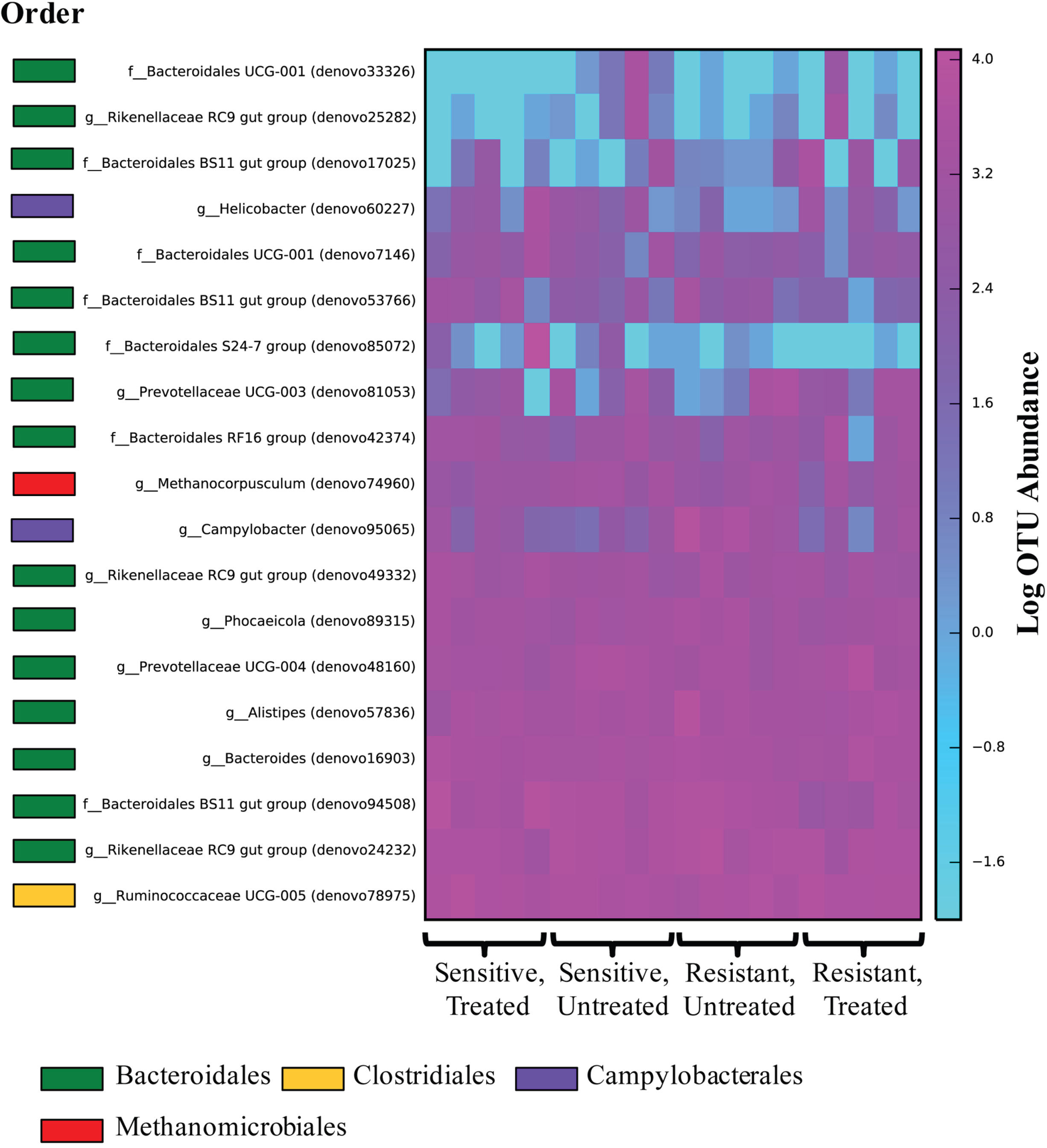
Heat map of abundances of individual OTUs in RF samples from monepantel-treated and untreated sheep, infected with sensitive or resistant *T. circumcincta* (n=5 for each treatment). The five most abundant OTUs in each sample were included in the analysis. The most accurate levels of taxonomic classification for each OTU is shown next to the OTU identifier, and Phylum membership is indicated with coloured bars.

LEfSe analyses (28) at the Genus level were therefore employed in order to detect genera elevated specifically in both sets of samples from those animals either infected or uninfected with *T. circumcincta*. In the RF samples (Fig. 6A), three genera were identified as significant biomarkers for infected sheep (groups SUT, RUT and RT), along with the Families *Microbacteriaceae* and *Mycoplasmataceae*, while in the uninfected sheep (group ST), eleven different taxa including three genera from the family *Ruminococcaceae*, the Order *Myxococcales* and the Genera *Aeriscardovia* and *Bifidobacterium* were significantly elevated (LDA score > 2.5). The p values of these associations varied between 0.001 and 0.05 (Table S5). When the data from the TIMS samples were analysed (Fig. 6B), a larger set of taxa were identified as biomarkers for infected sheep (genera including *Rikenellaceae* RC9, *Treponema* 2, *Prevotella* 1 and *Fibrobacter*, plus the entire Class *Deltaproteobacteria*), while the Genus *Sharpea*, the Family *Bifidobacteriaceae* and the Genus *Acetitomaculum* were identified as biomarkers for the uninfected group (LDA score >3.5). The p values of these associations varied between 0.0002 and 0.05 (Table S5). This suggests that, while the taxa which drive the differences seen at the Genus level between uninfected and infected sheep differ depending on whether RF or TIMS samples are analysed, specific elevation of several biomarkers is seen in uninfected animals at each sampling site, including sequences derived from the *Bifidobacteriaceae* at both sampling sites.

## DISCUSSION

Few studies have examined the host-microbiota-helminth relationship in animal hosts. To address this knowledge gap, the primary aim of this study was to investigate the species compositional changes in intestinal microbiota resulting from *T. circumcincta* infection and subsequent anthelmintic administration in sheep. However, it is well documented that factors such as age, diet, environment and sampling site are significant sources of inter-animal variation in microbial composition (29). To investigate this inherent variation between sheep, a preliminary trial was undertaken on the microbiota of the terminal ileum and faeces derived from age-matched lambs that were all grazed under similar conditions. By comparing the inherent variation between RF samples and TIC samples, our results suggest that there was significantly less variability of the microbiota between identically treated sheep in RF than in samples collected from the terminal ileum. Further, results showed that there were significant differences in microbial diversity between the RF and TIC samples with a reduction in alpha diversity in the latter. These findings were consistent with previously published results which showed that bacterial abundance in the large intestine was greater than in the small intestine and that the microbiota within the lumen of the terminal ileum is less diverse than in the rectum (29, 30). Bacterial taxonomic distribution patterns were also different between TIC and RF samples. *Clostridiales* was the dominant Order for all but one sample in the TIC, whereas the *Clostridiales* and *Bacteroidales* shared the dominant position in RF samples (Fig S1).

Although faeces are not necessarily the optimal correlate for examining the microbiota within a specific zone in the intestine (29, 31, 32), faecal samples were selected for the main study (to identify compositional differences of microbial communities between treated and untreated sheep, infected with resistant or sensitive *T. circumcincta*) due to the more consistent sample microbiota composition within each treatment group. In this study, the successful treatment of the infection did appear to have a significant impact on the diversity and composition of the faecal microbiota. The diversity of the microbiota of sheep successfully treated (ST) was significantly lower than that of the three groups of sheep that maintained the parasitic infection (SUT, RT, RUT). There are also compositional differences in the faecal microbiota of successfully-treated sheep, which are strongly correlated with a reduction in worm burden (as expected), and an increase in the abundance of the Phylum *Actinobacteria*. However, there was little difference in the diversity and abundance of the commensal microorganisms between the untreated sheep infected with the two different strains (resistant and sensitive) of parasitic nematode in the absence of treatment.

As a result of the large inter-sample variation between samples from the TIC (when compared to RF samples), the experimental design was modified to include the TIMS from nematode-infected sheep. This addition to the experimental design expanded the study to include an assessment of epithelial-attached microbiota in comparison to RF microbiota. The results derived from these latter samples should be taken with caution, as they required a different DNA extraction technique and the microbial DNA yield was much lower compared to the other samples in this study. However, the results of the 16S rDNA sequencing suggested that the samples were of low bacterial diversity and of very different composition from the faecal samples, with large inter-sample variation (similar to the TIC samples). This was expected as the resident gut microbes are known to be less abundant than and taxonomically-distinct from those which pass through the gut (29). Interestingly, microbial alpha-diversity of samples derived from the TIMS of the ST and RT groups was significantly decreased compared to those from the SUT and RUT groups, suggesting that in this location, MPTL treatment affects microbial diversity irrespective of whether the parasite infection is successfully treated or not. However, compositional (beta) diversity was affected by whether the infection was maintained, and these differences were strongly correlated to worm burden, a result consistent with that obtained from the RF samples. Unlike in faeces, where worm burden was strongly negatively correlated with the abundance of *Actinobacteria*, the abundances of *Bacteroidetes* and *Fibrobacteres* were positively correlated with worm burden in TIMS samples, while a negative correlation with the abundance of *Firmicutes* was observed.

Although different sets of biomarkers for parasite-infected and uninfected sheep were identified between the TIMS and RF datasets, some biomarkers were consistently associated with both sets of data. Species from the physiologically-uncharacterised *Rikenellaceae* RC9 gut group, which are abundant in rumen and faecal samples from other mammals (25, 33), were consistently elevated in both TIMS and RF samples from infected sheep. OTUs mapping to this Genus were also abundant in both sample sets (Figs. 5 and S4). Genera within the *Prevotellaceae*, known to be abundant in the rumen of sheep (34), were also biomarkers for nematode infection in both datasets. In contrast, members of the *Bifidobacteriaceae* were elevated in both TIMS and RF (represented by the genera *Aeriscardovia* and *Bifidobacterium*) of successfully treated sheep (ST). In the ileal epithelium, a single *Bifidobacterium* OTU (denovo3328) is clearly more abundant in the absence of *T. circumcincta* infection (Fig. S4). *Bifidobacteria* are abundant members of the gastrointestinal tract in mammals, and are thought to be acquired via maternal transmission in milk (35), although they have not previously been studied in the gastrointestinal tract of sheep. In humans, they are generally thought to be a marker of good gut health (35). It is also noteworthy that *Sharpea* species identified as the most significant biomarker for uninfected sheep in TIMS samples, also comprised one of the most abundant OTUs in these samples (denovo4171; Fig. S4). *Sharpea* has been identified as a Genus enriched in the rumen of low-methane yield sheep, where it is thought to be associated with rapid fermentation (36).

Although the direct effects of MPTL on the microbiota of sheep are not well characterised, in this study we were unable to detect differences in the RF samples due to MPTL application *perv se*. However, there was a small but significant decrease in diversity of the microbiota in TIMS samples from sheep treated with monepantel compared to the untreated sheep, and a compositional effect additional to that caused by the successful eradication of helminths. This effect may be a direct effect on the microbial community but, as with other anthelmintics, the effect may be immunological and/or physiological (37).

There is some inconsistency in the literature regarding the effect of helminth infection on microbial alpha-diversity in the gut. The increase in microbial diversity described here is consistent with a study of helminth-colonised humans in Malaysia (38), whereas controlled studies of *Trichuris muris* infection in mice (39, 40) suggest an association between helminth infection and decreased microbial diversity. There is also divergence between these studies and our results in terms of the beta-diversity changes associated with helminth infection and clearance, which are probably due to the different mammalian hosts, parasite species and conditions (natural versus experimental) studied. In the Malaysian human study, helminth infection was associated with an increase in *Paraprevotellaceae*, especially when *Trichuris* spp. were present (38). In contrast, the experimental mouse studies with controlled and treated *T. muris* infection both detected a reduction in members of the *Prevotellaceae* associated with infection, and in one case suggested transient elevation of *Bifidobacterium*, followed by *Lactobacillus*, in infected animals (39, 40). Treatment and clearance of the parasitic infection with the drug mebendazole enabled partial restoration of microbial alpha-and beta-diversity in infected animals, providing evidence that the parasitic infection was responsible for the induction and maintenance of the altered microbiota (39). While universal changes in richness and taxonomic composition due to helminth infection across multiple hosts and parasite species are unlikely to be found, it is established as a result of these previous studies and our work that these parasites cause changes in the gut microbiota in murine, ovine and human systems, which can be resolved by drug treatment.

In conclusion, we find microbiota variation in the ovine gut to be niche specific, as the microbiota sampled from either TIC or TIMS samples not only have reduced richness but are also much more variable between individual sheep when compared to the RF sample. However, in this study biomarkers were identified common to both the TIMS samples and the RF samples distinguishing sheep with helminth infection from those that cleared infection using anthelmintic treatment. In future studies, longitudinal investigation of changes in the microbiota of artificially or naturally infected animals with parasites and/or pathogenic bacteria will help clarify the causal relationship between infection, treatment and microbial composition. However, it is important to be aware that other factors may also play a part in alterations of the microbiota, including the effect of infection on metabolites (41). Further work will consider how the introduction of helminths can also alter the metabolome profiles within the intestine and the effect on the immune system (3, 42).

Understanding the dynamic mechanisms required to sustain and control the balance between pathogenic, beneficial and commensal microbial communities within the gastrointestinal tract will enable progress in the discovery and development of therapeutic reagents based on beneficial microorganisms and/or their excreted/secreted products to restore dysfunctional microbiotas or prevent the destabilisation of the microbiota by invading pathogens or opportunistic commensal bacteria (43, 44). These novel microbiota-based treatments will be particularly relevant to the problems of multidrug and anthelmintic resistance.

## MATERIALS AND METHODS

### Inherent variation of the gut microbiota between identically treated sheep from two different sampling areas of the gastrointestinal tract - the terminal ileum and rectum

Six one-year-old parasite-naïve lambs were grazed for four weeks on a paddock infected with a mixed population of ovine nematodes including *T. circumcinta* and *Haemonchus contortus*. Following grazing the lambs were re-housed for four weeks and necropsied as per (45). At post mortem faeces (1-2 g) were taken from the rectum, as well as the TIC. Samples were transferred to ice and stored at −80°C before processing.

### Effect of treatment on sensitive and resistant parasitic nematodes on the gut microbiota

The experimental design is outlined in (46). In brief, twenty parasite-naïve lambs of 8-9 months old were raised in containment, free from adventitious nematode contamination. These sheep were divided into two groups of ten animals and experimentally challenged *per os* with 7,000 *T. circumcincta* infective larvae (L_3_) of either a monepantel (MPTL)-sensitive (S) parental strain (designated MTci7) or an MPTL-resistant (R) strain that had been artificially derived from the parent strain (designated MTci7-12; Table 1).

Twenty-eight days post-infection the two groups were subdivided into groups of five animals and either left untreated (UT) to act as controls, or weighed and dosed orally by syringe at the manufacturer’s recommended dose rate with MPTL (2.5 mg kg^-1^ body weight; T). The four groups MTci7 untreated, MTci7 MPTL-treated MTci7-12 untreated and MTci7-12 MPTL-treated are designated as SUT, ST, RUT and RT respectively (Table 1). All of the animals were necropsied seven days post-treatment. Abomasa were collected and processed for worm burden estimation (45). Total worm burdens were used to confirm treatment outcome (Table 2).

**TABLE 2.** Sheep infection and treatment groups

| Sheep ID | Group ID | Nematode Isolate | Treatment | Faecal egg count (eggs g<sup>-1</sup> faeces) |  |
| --- | --- | --- | --- | --- | --- |
| OV1 | ST | Sensitive | MPTL | 0 |  |
| OV2 | ST | Sensitive | MPTL | 1 |  |
| OV3 | ST | Sensitive | MPTL | 0 | 777 |
| OV4 | ST | Sensitive | MPTL | 0 | 778 |
| OV5 | ST | Sensitive | MPTL | 0 |  |
| OV6 | SuT | Sensitive | Untreated | 360 | 779 |
| OV7 | SuT | Sensitive | Untreated | 45 | 780 |
| OV8 | SuT | Sensitive | Untreated | 33 |  |
| OV9 | SuT | Sensitive | Untreated | 63 | 781 |
| OV10 | SuT | Sensitive | Untreated | 360 |  |
| OV16 | RT | Resistant | MPTL | 333 | 782 |
| OV17 | RT | Resistant | MPTL | 333 | 783 |
| OV18 | RT | Resistant | MPTL | 333 |  |
| OV19 | RT | Resistant | MPTL | 270 | 784 |
| OV20 | RT | Resistant | MPTL | 1962 |  |
| OV11 | RuT | Resistant | Untreated | 972 | 785 |
| OV12 | RuT | Resistant | Untreated | 351 | 786 |
| OV13 | RuT | Resistant | Untreated | 792 |  |
| OV14 | RuT | Resistant | Untreated | 306 | 787 |
| OV15 | RuT | Resistant | Untreated | 387 |  |

RF samples were collected and stored at −80°C before processing. TIMS samples were also collected from these sheep. Briefly, terminal ileum tissue was thawed on ice and opened longitudinally, to reveal the mucosal layer of the lumen. The lumen was washed with PBS to remove debris before the mucosal layer was scraped off using a microscope slide, transferred to a 50 ml tube and vortexed for 30 seconds. A volume of 400 μl of scraped sample was added to 4.60 ml of RNAlater^®^ and stored at −80°C.

All experimental procedures described were approved by the Moredun Research Institute Experiments and Ethics Committee and performed under the legislation of a UK Home Office License (reference PPL 60/03899), complying with the Animals (Scientific Procedures) Act 1986.

### DNA extraction

Microbial genomic DNA was purified with a MO BIO PowerFecal™ DNA Isolation Kit, according to the manufacturer’s protocols. Briefly, 0.25 g of faeces or 0.25 ml of TIC were homogenized, using bead beating, to facilitate microbial cell lysis. The total microbial genomic DNA was eluted in DNA elution buffer. Additionally, DNA was extracted from TIMS samples using the Qiagen Tissue/Blood kit following the manufacturer’s protocol after bead beating as described above. Genomic DNA was quantified using NanoDrop™ spectrophotometry.

### Amplification of bacterial 16S rRNA genes

The bacterial 16S rRNA gene V4 region was amplified for Illumina MiSeq sequencing via a barcoded-adapter based PCR approach (47). PCR reactions for the bacterial 16S rRNA gene V4 region contained 1× *Taq* buffer plus additional MgCl_2_ (final concentration of 2.5 mM), 0.2 mM of each of the four dNTPs, 0.25 μM of each primer, 0.05 U μl^-1^ *Taq* DNA polymerase, and 1 ng μl^-1^ template DNA in a total volume of 25 μl with PCR grade water, set up under contaminant-free conditions (48). For each amplification reaction the same forward primer (515F) together with a different barcoded reverse primer (806R) was used (the reverse primer sequences differed only at the barcode region (47)). Amplification of the bacterial 16S rRNA gene V4 region was as follows: 94°C for 3 min; followed by 25 cycles at 94°C for 45 sec, 50°C for 60 sec, 72°C for 90 sec; followed by a single cycle of 72°C for 10 min. For TIMS samples, which contained a low amount of target DNA, 35 cycles of amplification via the above protocol were used. Because of this extended amplification protocol, a PCR negative control showed the presence of amplified sequences corresponding to possible contamination: two such OTUs assigned to the genus *Staphylococcus* and the species *Bradyrhizobium elkanii* were later removed from the dataset as a result.

Size and concentrations of PCR amplicons were analysed by agarose gel electrophoresis. The PCR amplicons were gel-purified using a Wizard^®^ Gel and PCR Clean-Up System (Promega, UK) and quantified using a Quant-iT PicoGreen ds DNA Assay Kit (Life Technologies, UK) before pooling in equimolar quantities for Illumina sequencing.

### Preparation of PCR amplicons for Illumina sequencing

Pooled PCR amplicons were sequenced at Edinburgh Genomics, University of Edinburgh. Paired-end sequencing (2×250 bp) was run on the Illumina MiSeq platform and ~11M raw read clusters were generated. Three separate sequencing primers were used for sequencing; two used to read sequences from either end: Read 1 primer (TATGGTAATT GT GTGCCAGCMGCCGCGGTAA) to yield the 5’ read; Read 2 primer (AGTCAGTCAG CC GGACTACHVGGGTWTCTAAT) to yield the 3’ read and a third as the indexing primer (ATTAGAWACCCBDGTAGTCCGGCTGACTGACT) used to read the barcode sequence (47).

### Data processing and filtering

Raw Illumina sequence reads were analysed using the QIIME workflow (49). Demultiplexed forward and reverse sequence reads were paired with a minimum overlap of 200 bp for maximum accuracy (50) before quality filtering with a minimum quality score of 20. As the bacterial 16S rRNA gene V4 region is universally bigger than 250 bp, short reads below this cut-off were removed using the Python script filter_short_reads.py (https://gist.github.com/walterst/7602058). Chimeric sequences were also removed using UCHIME (51) with version 128 of SILVA (52) as the reference database. Total numbers of quality-filtered sequences were ~2.4M for the inherent variation experiment (n = 2×6) and ~4.4M (RF) or ~446K (TIMS) for the nematode infection experiment (n = 4×5). *De novo* OTU picking at 97% similarity and taxonomic assignments against SILVA 128 were performed using Uclust; OTU representative sequences which failed to align with the dataset using PyNast (53) and singleton OTUs were removed before OTU tables were generated. OTU data were summarized by taxonomic ranks and via heat maps, and alpha rarefaction curves for observed species and Shannon diversity were generated using QIIME.

### Multivariate statistical analysis

Relative abundance OTU data were imported into Primer 6 Version 6.1.12 (Primer-E, Ivybridge, UK) and used to generate Bray-Curtis similarity matrices. PCoA and CAP ordinations were performed on the Bray-Curtis similarity matrices in Primer 6. Additional multivariate statistical analyses, PERMANOVA and PERMDISP (27) were performed with the PERMANOVA+ add on package for Primer 6. PERMANOVA and PERMDISP were used to test for significant differences in the distribution and dispersion of sample groups based on Bray-Curtis similarities. Linear discriminant analysis effect size (LEfSe) analysis (28) was performed on relative abundance taxonomy tables generated in QIIME using the LEfSe online Galaxy tool (http://huttenhower.sph.harvard.edu/galaxy/).

Accession number: Sequencing data and metadata were uploaded to the European Nucleotide Archive (ENA) at the European Bioinformatics Institute (EBI); study accession number PRJEB24185. http://www.ebi.ac.uk/ena/data/view/PRJEB24185

## ACKNOWLEDGEMENTS

We gratefully acknowledge funding from The Scottish Government’s Rural and Environment Science and Analytical Services Division (RESAS) and the M.Sc. Biotechnology programme of the University of Edinburgh. B.Y. was supported by the Erasmus Programme, and M.J.R.S. was supported by the Xunta de Galicia under the Galeuropa programme. We are also grateful to the Bioservices Division, Moredun Research Institute, for expert care and assistance with animals.

---

**FIG 6** Bar graph representation of taxonomic biomarkers identified in RF samples (A) and TIMS (A) from worm-free sheep (open bars) or worm-infected sheep (solid bars). LDA scores of >2.5 in panel A and >3.5 panel B are shown.

**FIG S1** Percentage abundance plot of taxonomic orders in TIC and RF samples. Orders with abundances <4% in all samples were grouped as “Others”.

**FIG S2** Percentage abundance plot of taxonomic orders in RF samples from monepantel-treated and untreated sheep, infected with sensitive or resistant *T. circumcincta* (n=5 for each treatment). Orders with abundances <4% in all samples are groups as “Others”.

**FIG S3** (A) Alpha rarefaction curves for OTUs in TIMS from monepantel-treated and untreated sheep infected with sensitive or resistant *T. circumcincta* (n=5 for each treatment). Error bars represent standard deviations. (B) Percentage abundance plot of taxonomic orders in TIMS from monepantel-treated and untreated sheep, infected with sensitive or resistant *T. circumcincta* (n=5 for each treatment). Orders with abundances <4% in all samples are grouped as “Others”.

**FIG S4** Heat map of abundances of individual OTUs in TIMS samples from monepantel-treated and untreated sheep, infected with sensitive or resistant *T. circumcincta* (n=5 for each treatment). The 5 most abundant OTUs in each sample were included in the analysis. The most accurate levels of taxonomic classification for each OTU is shown next to the OTU identifier, and Phylum membership is indicated with coloured bars. Full details of the percentage abundance and taxonomy of these OTUs in each sample is presented in Table S4.

